# Inspiration of SARS-CoV-2 envelope protein mutations on pathogenicity of Omicron XBB

**DOI:** 10.1101/2023.01.09.523338

**Authors:** Yi Wang, Hongying Ji, Xiaoli Zuo, Bingqing Xia, Zhaobing Gao

**Author notes:** Corresponding authors (Z.G.), (B.X.).

## Abstract

Predicting pathogenicity of Omicron sub-variants is critical for assessing disease dynamics and developing public health strategies. As an important virulence factor, SARS-CoV-2 envelope protein (2-E) causes cell death and acute respiratory distress syndrome (ARDS)-like pathological damages. Evaluation of 2-E mutations might offer clues to pathogenicity forecast. Here, the frequency and cell lethality of 92 mutations of 2-E in five early “variants of concern” (VOCs, Alpha, Beta, Gamma, Delta, and Omicron BA.1, BA.2, BA.3, BA.4, and BA.5) were analyzed, which could be divided into three classes. Most (87) mutations belong to Class I, no obvious frequency changes. Class II consists of 2 mutations, exhibiting enhanced cell lethality but decreased frequency. The rest 3 mutations in Class III were characterized by attenuated cell lethality and increased frequency. Remarkably, the Class II mutations are always observed in the VOCs with high disease severity while the Class III mutations are highly conserved in the VOCs with weakened pathogenicity. For example, P71L, the most lethal mutation, dropped to nearly 0.00% in the milder Omicrons from 99.12% in Beta, while the less lethal mutation T9I, sharply increased to 99.70% in BA.1 and is highly conserved in BA.1-5. Accordingly, we proposed that some key 2-E mutations are pathogenicity markers of the virus. Notably, the highly contagious Omicron XBB retained T9I also. In addition, XBB gained a new dominant-negative mutation T11A with frequency 90.52%, exhibiting reduced cell lethality, cytokine induction and viral production capabilities *in vitro*, and particularly weakened lung damages in mice. No mutations with enhanced cell lethality were observed in XBB. These clues imply a further weakened pathogenicity of XBB among Omicron sub-variants.

## Text

The Omicron, a SARS-CoV-2 variant of concern (VOC), was first detected in South Africa in mid-November 2021 ^1^. There was a surge of new emergent Omicron variants since the restrictions that were used to quash the virus’s spread were dismantled. The advantage sub-variants such as BA.1, BA.2, BA.3, BA.4, BA.5 had distinct transmission, neutralization and immune escape capabilities ^2^. Mutations in the viral spike (S) protein have been demonstrated to be responsible for immune escape and enhanced transmission ^3–5^. In comparison with the original strain and other variants, the pathogenicity of Omicron variants was milder and the rate of hospitalization and mortality were significantly reduced ^6^. However, it is worth to note, BA.5 infection has shown increased rate of recovery positive, raised proportion of infections who were “symptomatic” and some patients regained pneumonia symptoms ^7–9^. These phenomena remind us to be alert to the change in pathogenicity. However, the determinants of SARS-CoV-2 pathogenicity remain elusive. On December 10th, Chinese government lifted its strict “zero COVID-19” measures, which ended a massive lockdown of entire cities and has led to infection waves. Large-scale and short-term infection might bring natural and endless “evolution” places for viruses. The genomes of the Omicron variants continue to diversify apparently, and it is urgent to predict and monitor ‘dangerous’ mutations of pathogenicity.

The envelope protein of SARS-CoV-2 (2-E) forms a homo pentameric cation channel that is important for virus virulence ^10^. One of our previous studies indicated that 2-E channel alone is sufficient to induce cell death, provoke cytokine storm and even cause acute respiratory distress syndrome (ARDS)-like damages *in vivo ^10^*. Furthermore, T9I, a single high-frequency mutation of 2-E protein in Omicrons, was identified to reduce the virus replication and virulence through altering channel function in another study ^11^. These above results supported the mutation of 2-E protein might affect the evolution trend of virus pathogenicity. To further understand the potential contribution of 2-E mutations to pathogenicity, we measured the cell lethality of the 2-E spontaneous mutations with a frequency ≥0.01% in five VOCs (Alpha, Beta, Gamma, Delta, and Omicron) up to October 2022, and analyzed the correlation between cell lethality, frequency and clinical severity in the current study.

Based on the National Genomics Data Center (NGDC), there are 92 2-E mutations with a frequency ≥0.01% in the five VOCs (Table 1, Fig.S1 a). Among these mutations, there are 4 mutations with frequency higher than 1.00% and 2 of them higher than 90.00%. Omicron retained 31 mutations that emerged from the early 4 VOCs, and gained 7 new mutations (Fig.S1 b-c). We defined the difference between the highest frequency value of each mutation in the early four VOCs (Alpha, Beta, Gamma, Delta) and the highest frequency of Omicron BA.1-5 as the frequency change (ΔFrequency). Among them, 13 mutations exhibited increased frequency while 71 mutations showed decreased frequency (Fig.1 a, Fig.S1 a). Then the cell lethality of all 92 mutations were measured. Refer to our previous work, 92 2-E mutants were expressed in Vero E6 cells and the cell lethality was calculated through the ratio of cell death rate to the protein expression level (Fig.1 b-c, Fig.S2 a-b). In comparison with wild-type (WT) 2-E protein, 13 mutations introduced stronger capability of killing cells, while 51 mutations attenuated the capability. The rest 28 mutations failed to affect the cell lethality significantly (Fig.1 c).

**Table.1.**
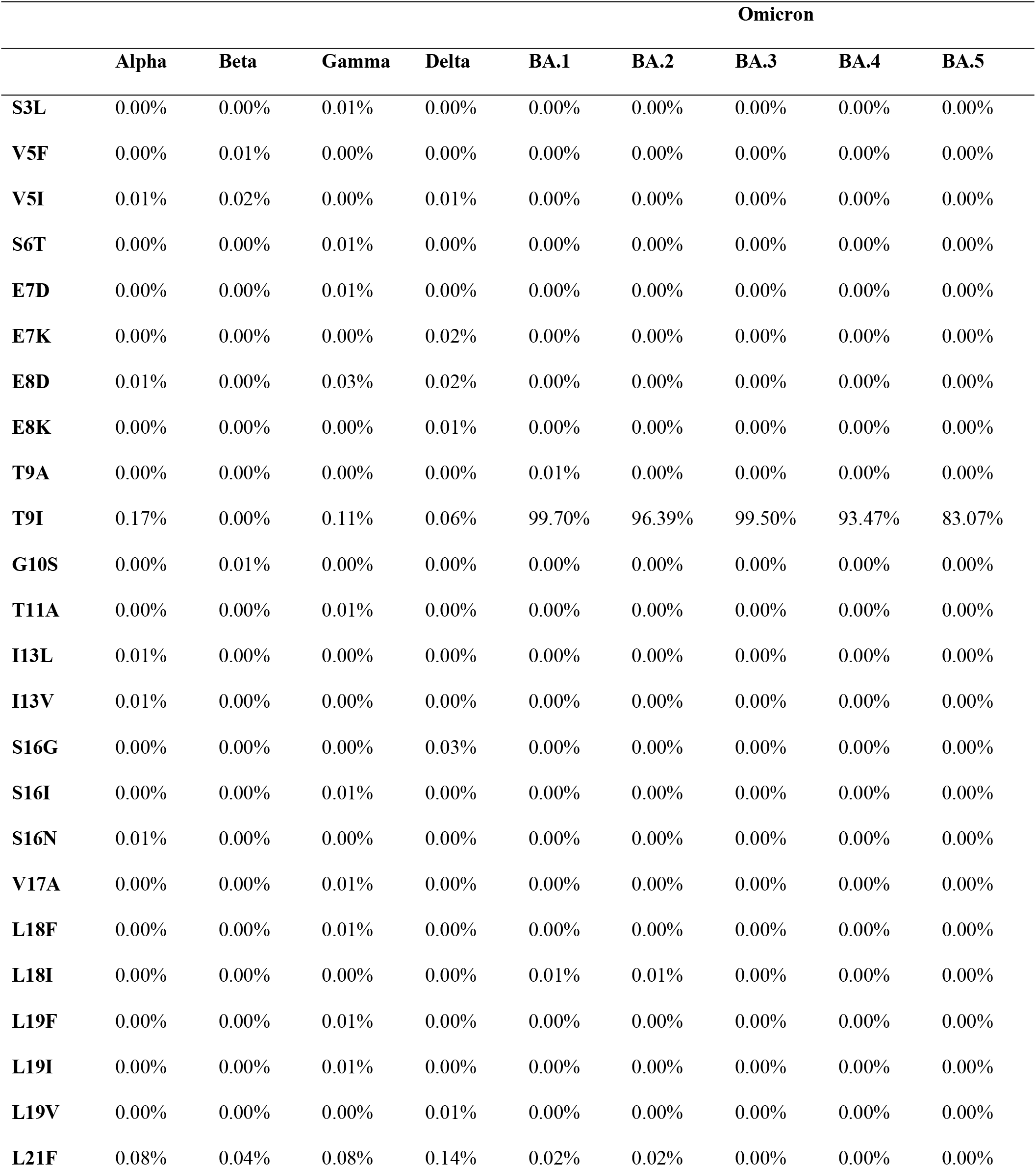
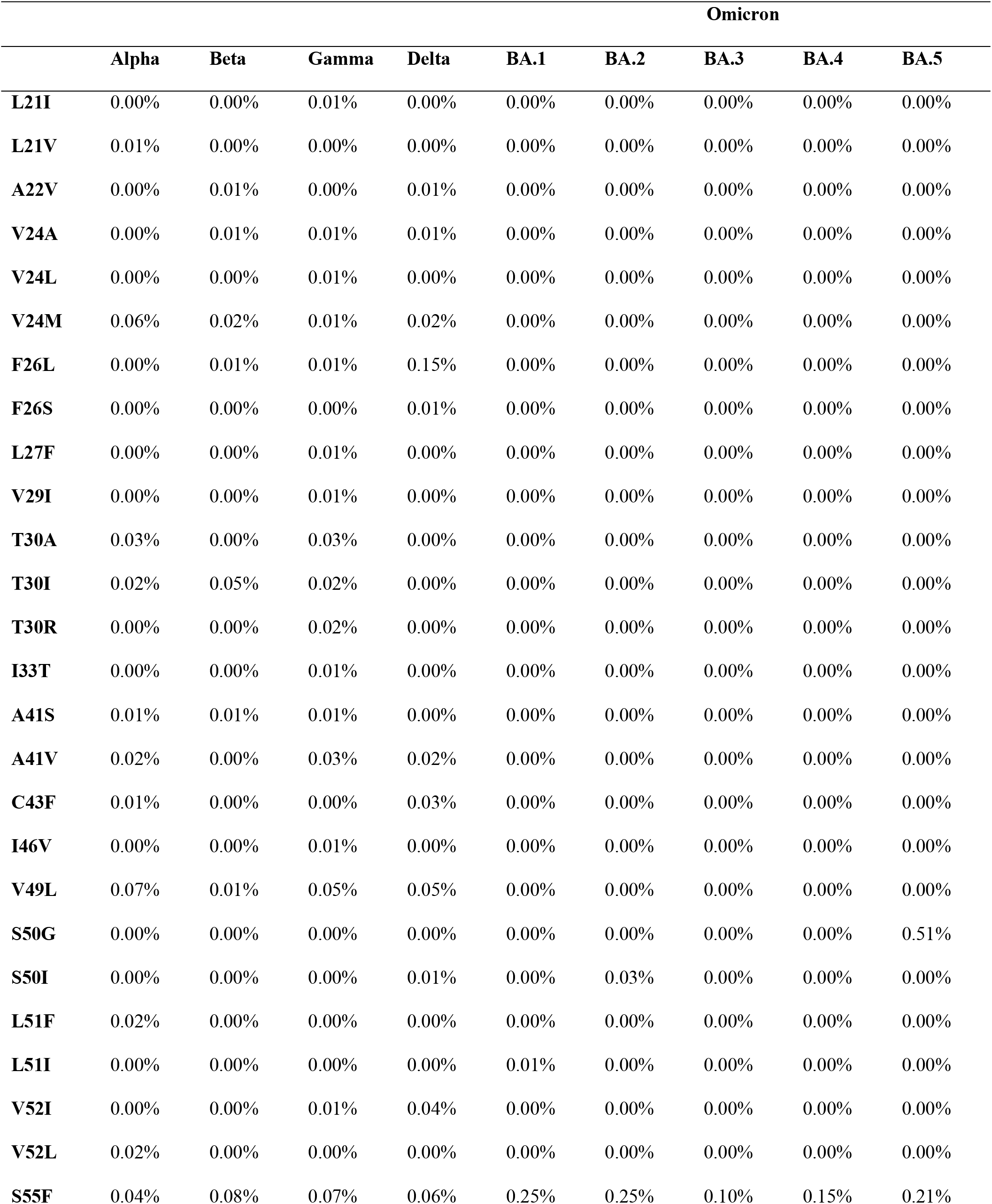
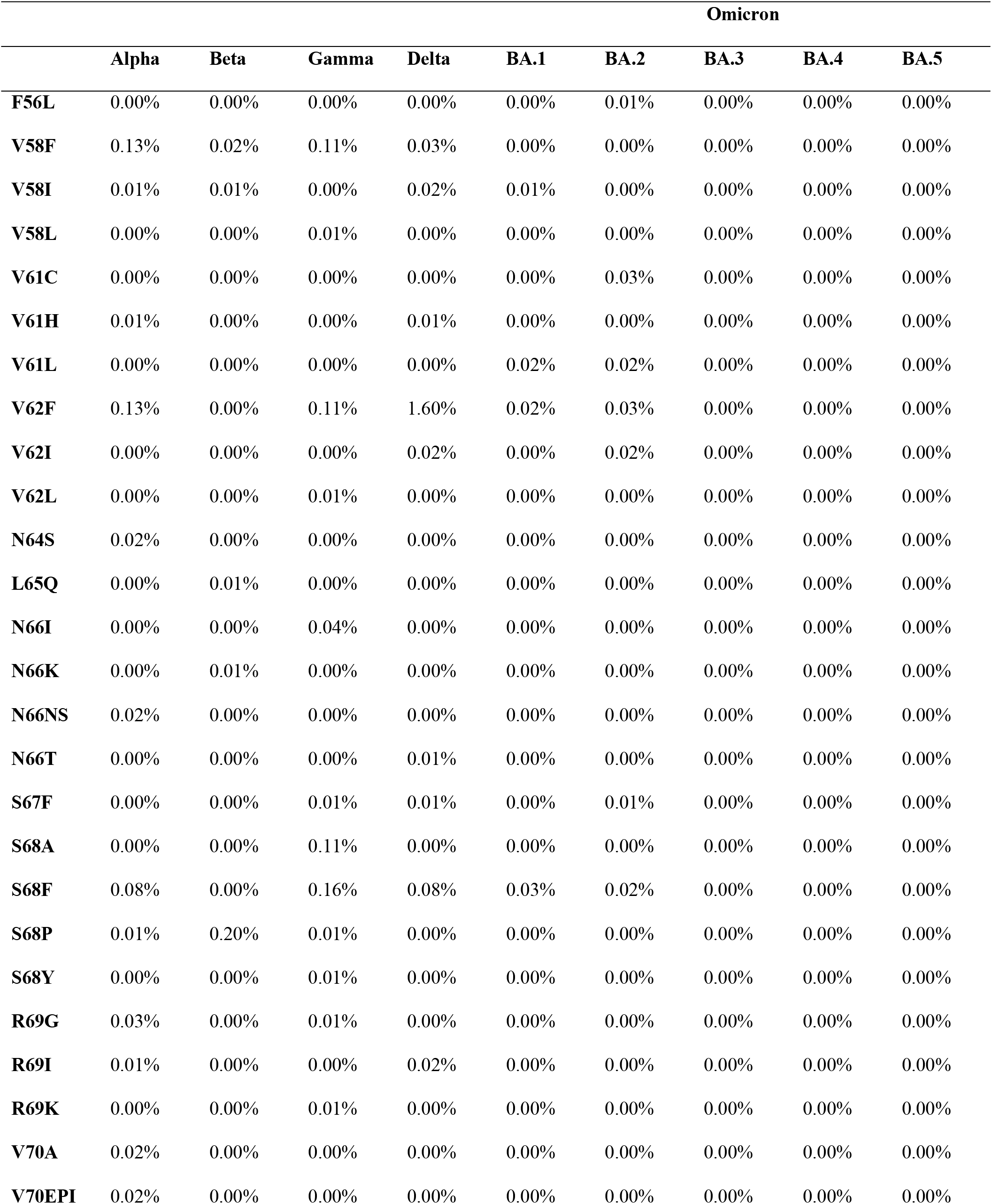
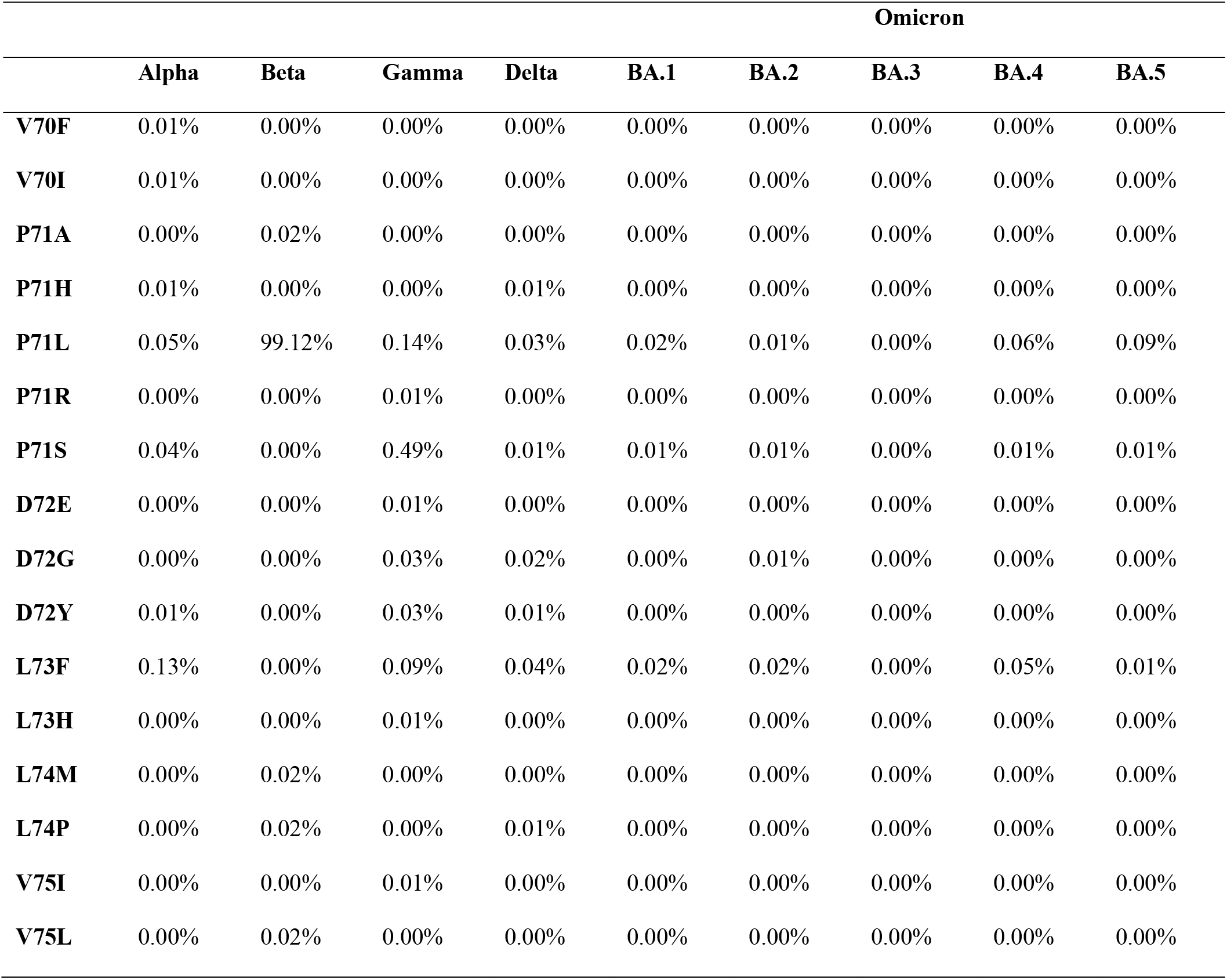
2-E mutations with frequency ≥ 0.01% in five VOC

Next, we analyzed the correlation between the cell lethality and *Δ*Frequency of each mutation. All mutations could be clearly distinguished into three groups, which were named as Class I, Class II, and Class III. Class I, the group contains 87 mutations with *Δ*Frequency ranging −0.20% to 0.20%. We speculated these mutations with low *Δ*Frequency are of less importance for virus phenotypes and failed to become advantage mutations. Interesting to note, all the 28 mutations that did not affect cell lethality belong to this class. Class II contains two mutations, P71L and P71S, which increase cell lethality significantly. The rest 3 mutations, S50G, S55F, and T9I, exhibiting increased frequency and reduced cell lethality, were classified into Class III. In general, the *Δ*Frequency seems was negatively correlated with the cell lethality (Fig. 1d), which inspired us to further explore the correlation between 2-E mutations and virus pathogenicity.

**Fig.1.**
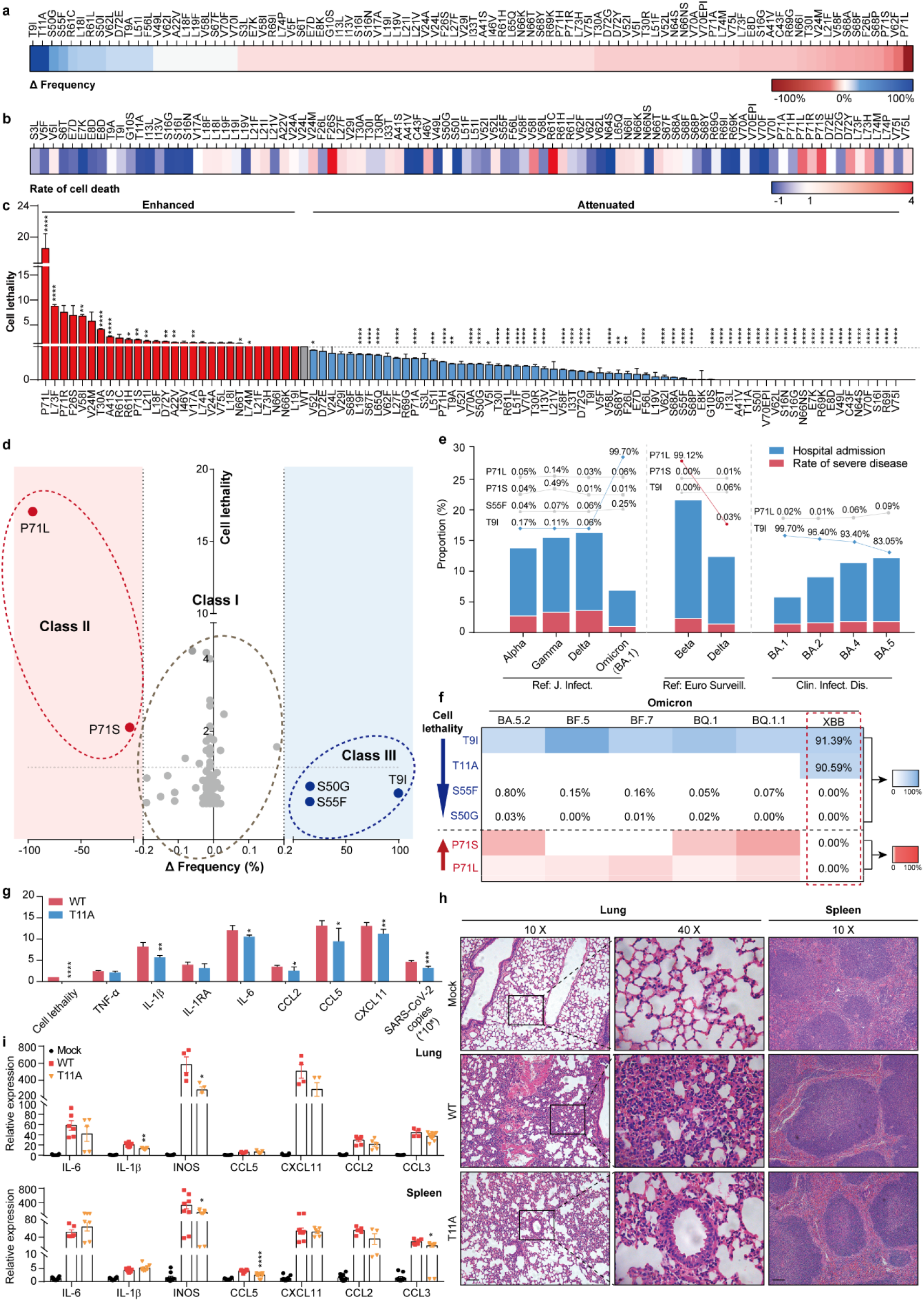
SARS-CoV-2 2-E mutations are potential pathogenicity markers. **a**, The Δ frequency of 92 2-E mutations. **b**, The rate of cell death of 2-E mutations. **c**, The cell lethality of 2-E mutations. The cell lethality was calculated through the ratio of cell death rate to the protein expression level. **d**, Correlation analysis of Δfrequency and cell lethality. Dotted gray circle represents Class I, dotted red circle represents Class II, dotted blue circle represents Class III. **e**, The quantification of hospitalization rate and disease severity up to Omicron BA.5 and tabulated the contribution of Class II and III mutations. Pathogenicity of different SARS-CoV-2 variants ^9, 12, 13^. **f**, Heat map of 6 key mutations frequency in Omicron sub-variants. **g**, The activity of 2-E WT and T11A in causing cell lethality, cytokine release and viral production. **h**, Histopathology of lungs and spleens from 2-E WT and T11A protein treatment groups (bar, 100 μm). **i**, qRT-PCR analysis of cytokine levels 6 h after treatment. *\*p < 0.05; **p <0.01; ***p < 0.001;* unpaired Student’s t test. All error bars are SEM (n≥3).

According to three independent clinical studies, we quantified the hospitalization rate and disease severity up to Omicron BA.5, and tabulated the contribution of the above listed mutations. The results suggest the mutations in Class II and III are perhaps the essential factors for disease severity. First, Class III (less lethal) mutations appeared more often in the milder variants than that in severe variants (Fig.1 e left). The most representative mutation in this class is T9I, sharply increased to 99.70% in milder BA.1 and is highly conserved in Omicron BA.1-5 (Table 1). In contrast, its frequency in more severe variants such as Alpha, Gamma and Delta are as low as 0.17%, 0.11% and 0.06%, respectively. Interestingly to note, the frequency of S55F, another less lethal mutation, increased around 4-times in Omicron BA.1 also^12^. Second, Class II (more lethal) mutations seems are correlated to more severe variants (Fig.1e, middle). Mutation P71L appeared in Alpha and the frequency raised to 99.12% in Beta, the most severe variant so far^13^. After the emergence of Omicrons, the frequency of P71L decreased sharply or even disappeared in BA.3 (Table 1). As for P71S, it emerged in Alpha, reached its highest frequency in Gamma (0.49%), and gradually disappeared in Omicron. Third, lightly frequency change of T9I (Class II) and P71L (Class III) mutations may affect the virus pathogenicity particularly among milder Omicron sub-variants (Fig.1 e, right). Compared with Omicron BA.1 and BA.2, the clinical symptoms of BA.4 and BA.5 are more severe^9^. Correspondingly, the frequency of T9I dropped 4.30% and 16.65% in the latter two sub-variants, respectively. The more lethal P71L increased gradually and reached to 0.09% in BA.5, its third highest frequency among the monitored variants. These results highlighted the important roles of 2-E mutations in determining virus pathogenicity. We proposed the five mutations in Class II (P71L and P71S) and III (T9I, S55F and S50G) may act as pathogenicity markers of SARS-CoV-2.

Following BA.5, various new sub-variants appeared in Omicron, including BA.5.2 and BF.7, two major pandemic variants in China at present. We then supervised the five potential pathogenicity markers in the latest six sub-variants before December 2022. We found that T9I mutation is still conserved in these sub-variants, e.g. BA.5.2 (91.68%) and BF.7 (91.44%). The frequency changes of S55F and P71L in BA5.2 and BF.7 drew our attention also. In BA5.2, the frequency of S55F increased to 0.80% and that of P71L decreased to 0.02%. In BF.7, the frequency of S55F lightly decreased to 0.15% while that of P71L increased to 0.05%. On December 20, 2022, the Chinese Center for Disease Control and Prevention announced that XBB is a new variant branch of Omicron that has been imported into China. Although XBB is known as the “strongest immune escape variant”, its pathogenicity remains unclear ^14^. Encouragingly, we found that XBB retained the mutation T9I and notably, gained a new mutation T11A, and the frequency reached to 90.50%. One of our previous studies has demonstrated that T11A is a dominant-negative mutation of channel function ^10^. Besides impairing the channel activity, in comparison with WT 2-E protein, T11A expression significantly alleviated cell death and caused less cytokine release. In addition, the capability of virus releasing and virulence were also weakened (Ref. Fig.1 g). In this study, the influence of T11A was further evaluated *in vivo*. In comparison with WT proteins, inoculation T11A proteins caused weaker lung pathological phenotype in mice (Fig.1 h). We observed marked inflammatory cell infiltration, edema, pulmonary interstitial hyperemia, hemorrhage, and alveolar collapse in WT group. In contrast, severe damages in T11A and buffer solution groups were not observed (Fig.1 h). The cytokines were further characterized using qRT-PCR. In comparison with WT group, the expression levels of cytokines (IL-1β, IL-6, and INOS, etc.) and chemokines (INOS, CCL3 and CCL5, etc.) were much lower in T11A groups (Fig.1 i). These clues imply a further weakened pathogenicity of XBB sub-variant.

Omicron is continuously evolving. Predicting and rapidly characterizing virus pathogenicity of new variants is critical for assessing disease dynamics and developing public health strategies. In this study, we found that the frequency change and cell lethality of some 2-E mutations might be determinants of virus pathogenicity. Accordingly, five 2-E mutations were proposed to be potential pathogenicity markers. We applied our predictive theoretical model to forecast the potential pathogenicity of XBB. Two high-frequency mutations T9I and T11A with reduced cell lethality were observed, which might confer weaker pathogenicity to XBB. Nevertheless, we still need to be vigilant whether there will be a sudden frequency increase of highly pathogenic mutations such as P71L or the emergence of new 2-E mutations in the future. Although the existing literature on SARS-CoV-2 transmission, immune escape and evolutionary analysis is vast, we believe this study is the only one that systematically links 2-E mutations to virus pathogenicity. The analyses performed here come with their limitations. First, the influence of the proposed mutations needs to be verified at virus level. Second, the data linking genomic sequencing results of 2-E and clinical patient severity is deficient. Nevertheless, as some countries are at the peak of infections, our findings might provide advance warning of potential outbreaks.

## Acknowledgement

We are grateful to the National Science Fund of Distinguished Young Scholars (81825021), Fund of Youth Innovation Promotion Association (2019285), the National Natural Science Foundation of China (31700732, 81773707, 92169202), the Linggang lab (LG202101-01-04), Fund of Shanghai Science and Technology (20ZR1474200, 22QA1411000).

## Author contributions

Z. G., conceived designed the project. B.X., Y.W., H.J. and X.Z. carried out the cell-based assays; B.X., Y.W. and X.Z. carried out the animal experiments with purified proteins. Z. G., B. X. wrote the manuscript. All authors read and approved the manuscript.

## Methods

### Mutation frequency statistics

The frequency of 2-E spontaneous mutations was collected from National Genomics Data Center (NGDC). All mutations with frequency ≥ 0.01% were selected to further analysis. The Δfrequency was defined as difference of the maximum frequency value of early VOCs 2-E mutations and that of Omicron sub-variants.

### Plasmids and mutagenesis

Wild-type SARS-CoV-2-E sequences were synthesized in pcDNA3.1 by the Beijing Genomics Institute (BGI, China). Point mutations were generated using sited-directed mutagenesis, and all mutations were confirmed by sequencing (BGI, China).

### Cell culture

Vero E6 cells were purchased from National Collection of Authenticate Cell Cultures (China). Vero E6 cells were grown in 90% DMEM basal medium (Gibco, NY, USA) supplemented with 10% fetal bovine serum (Gibco, NY, USA) and 100 units/mL penicillin/streptomycin (Gibco, NY, USA). Cells were grown at 37 °C, 5% CO2 incubator, and passaged approximately every 2 days when on fluency up to 80%–90%. Vero-E6 cells were seeded on the 6-well cell culture plates for western blot and 96-well culture plates for CCK-8 assay.

### Western blot

Cells were lysed by lysis buffer (20118ES60, Yeasen, China). Then the proteins were collected and quantified by BCA protein assay kit (Thermofisher, USA). Proteins were resolved in 12% SDS-PAGE, transferred to PVDF membranes (GE, USA), and incubated with primary antibodies against HA-Tag (proteintech, USA), GAPDH (Yeasen, China). Second antibodies are peroxidase-Conjugated Goat Anti-Rabbit IgG (H+L) (33101ES60, Yeasen, China) and Peroxidase AffiniPure Goat Anti-Mouse IgG (H+L) (33201ES60, Yeasen, China). Then the relative expressions of 2-E mutations were analyzed with ImageJ.

### Relative cytotoxicity assay

Vero E6 cells were transfected with 400 ng/well 2-E plasmids or 2-E mutations using Lipofectamine 3000 Transfection Reagent (L3000015, Thermo Fisher, MA, USA). The control group was Vero E6 cells transfected with 400 ng/well pcDNA3.1 vector plasmids alone. After 24 h, we tested cell viability by CCK-8 kit (40203ES60, Yeasen, China) according to our previous work. Absorbance analysis were performed with Thermo Scientific Microplate Reader at 450 nm according to the manufacturer’s instructions (Thermo Fisher, MA, USA). Normalized cytotoxicity = (Ablank – Amutaion)/(Ablank– Amodel) %.

### Animal model

2-E WT or T11A protein was conducted tail vein injection in 8-week-old mice (25 mg/kg body weight) to establish mouse model. Male C57BL/6 mice were obtained from Shanghai SLAC Laboratory Animal Co., Ltd. All animal procedures were performed in accordance with the National Institutes of Health Guide for the Care and Use of Laboratory Animals, under protocols approved and strictly followed by the Institutional Animal Care and Use Committees (IACUC).

### Histology

Tissues from 2-E WT or T11A treated mice were fixed in 4% PFA for at least 7 days, and then were paraffin embedded. The paraffin samples were cut into 3-μm sections following the standard procedure. The sections were stained with H&E, and examined by Leica TCS-SP8 STED system (Leica Microsystems, DE).

### qRT-PCR Analysis

Total RNAs were extracted from cells and tissues using Trizol (Invitrogen, USA) and all total Nucleic Acid Isolation Kit (Ambion Inc., USA), following the manufacturer’s instruction. The experiment was performed using SYBR Green Master Mix (11184ES03, Yeasen, China) to quantify the mean values of delta Ct and SEM (n≥ 3). The primers used for quantification were listed in Supplementary information, Table S1.

---

**Table.2.** 2-E mutations with frequency ≥ 0.01% in newly emergent Omicron sub-variants

|  | **BA.5.2** | **BF.5** | **BF.7** | **BQ.1** | **BQ.1.1** | **XBB** |
| --- | --- | --- | --- | --- | --- | --- |
| **V5I** | 0.01% | 0.01% | 0.11% | 0.01% | 0.01% | 0.00% |
| **E7K** | 0.01% | 0.01% | 0.00% | 0.00% | 0.00% | 0.00% |
| **E8D** | 0.00% | 0.00% | 0.00% | 0.00% | 0.01% | 0.00% |
| **T9A** | 0.01% | 0.01% | 0.00% | 0.01% | 0.00% | 0.00% |
| **T9I** | 91.68% | 96.76% | 91.44% | 93.13% | 91.19% | 91.39% |
| **T11A** | 0.00% | 0.00% | 0.00% | 0.00% | 0.01% | 90.59% |
| **S16G** | 0.00% | 0.00% | 0.00% | 0.00% | 0.01% | 0.00% |
| **L18I** | 0.04% | 0.01% | 0.05% | 0.00% | 0.00% | 0.00% |
| **L21F** | 0.03% | 0.01% | 0.03% | 0.06% | 0.02% | 0.00% |
| **L21I** | 0.00% | 0.00% | 0.03% | 0.00% | 0.00% | 0.00% |
| **A22V** | 0.00% | 0.00% | 0.00% | 0.01% | 0.00% | 0.00% |
| **V24L** | 0.01% | 0.00% | 0.00% | 0.00% | 0.00% | 0.00% |
| **V24M** | 0.01% | 0.00% | 0.00% | 0.02% | 0.01% | 0.00% |
| **F26L** | 0.01% | 0.00% | 0.00% | 0.01% | 0.00% | 0.00% |
| **T30I** | 0.00% | 0.00% | 0.00% | 0.00% | 0.01% | 0.00% |
| **A41V** | 0.01% | 0.00% | 0.00% | 0.01% | 0.00% | 0.00% |
| **C43F** | 0.00% | 0.00% | 0.00% | 0.01% | 0.00% | 0.00% |
| **V49L** | 0.01% | 0.01% | 0.00% | 0.00% | 0.00% | 0.00% |
| **S50G** | 0.03% | 0.00% | 0.01% | 0.02% | 0.00% | 0.00% |
| **S50I** | 0.01% | 0.05% | 0.00% | 0.00% | 0.00% | 0.00% |
| **L51F** | 0.00% | 0.00% | 0.00% | 0.00% | 0.00% | 0.00% |
| **L51I** | 0.01% | 0.01% | 0.00% | 0.00% | 0.02% | 0.00% |
| **S55F** | 0.80% | 0.15% | 0.16% | 0.79% | 0.05% | 0.07% |
| **V58F** | 0.01% | 0.00% | 0.00% | 0.00% | 0.00% | 0.00% |
| **V58I** | 0.01% | 0.03% | 0.00% | 0.01% | 0.01% | 0.00% |
| **V58L** | 0.02% | 0.00% | 0.00% | 0.00% | 0.02% | 0.00% |
| **R61C** | 0.01% | 0.00% | 0.00% | 0.00% | 0.00% | 0.00% |
| **R61L** | 0.01% | 0.00% | 0.19% | 0.00% | 0.00% | 0.00% |
| **V62F** | 0.27% | 0.07% | 0.21% | 0.01% | 0.06% | 0.00% |
| **V62I** | 0.01% | 0.04% | 0.00% | 0.00% | 0.00% | 0.00% |
| **N66K** | 0.00% | 0.01% | 0.01% | 0.00% | 0.00% | 0.00% |
| **S68F** | 0.04% | 0.06% | 0.00% | 0.02% | 0.00% | 0.00% |
| **S68P** | 0.00% | 0.02% | 0.02% | 0.01% | 0.00% | 0.00% |
| **V70A** | 0.00% | 0.00% | 0.01% | 0.00% | 0.00% | 0.00% |
| **V70I** | 0.00% | 0.00% | 0.00% | 0.00% | 0.01% | 0.00% |
| **P71H** | 0.01% | 0.00% | 0.01% | 0.00% | 0.00% | 0.00% |
| **P71L** | 0.02% | 0.03% | 0.05% | 0.02% | 0.04% | 0.00% |
| **P71S** | 0.09% | 0.00% | 0.00% | 0.07% | 0.11% | 0.00% |
| **D72G** | 0.01% | 0.13% | 0.01% | 0.00% | 0.01% | 0.00% |
| **L73F** | 0.01% | 0.02% | 0.05% | 0.01% | 0.01% | 0.10% |

**Table S1:**
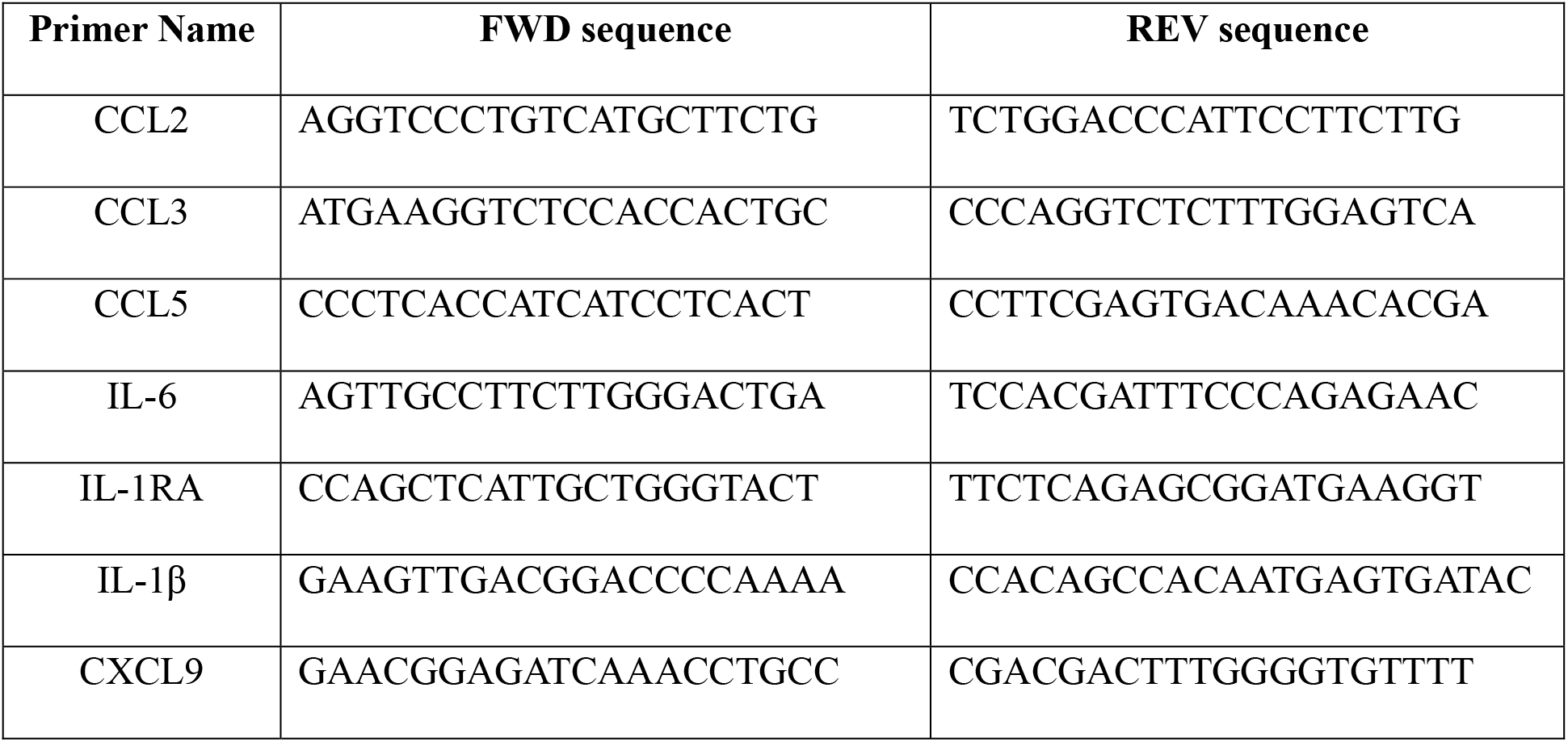
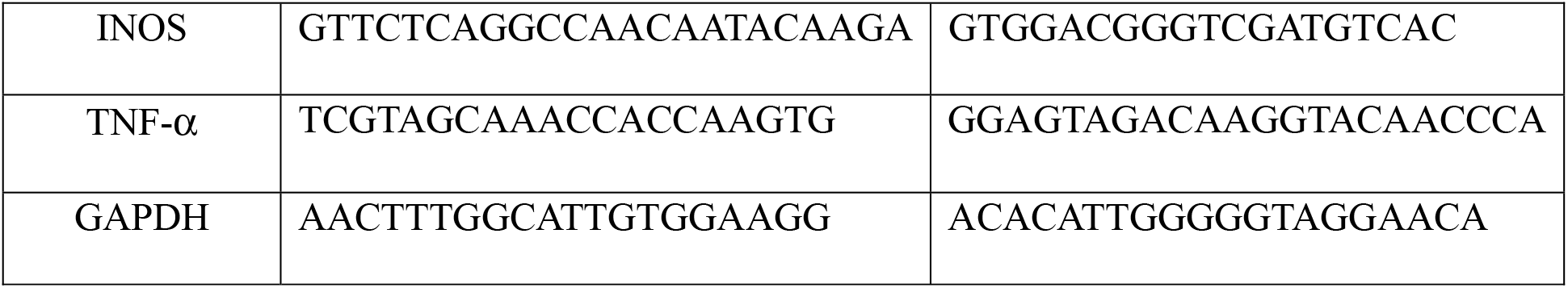
List of Quantitative Real-time PCR (qRT-PCR) primers.

**Fig.S1.**
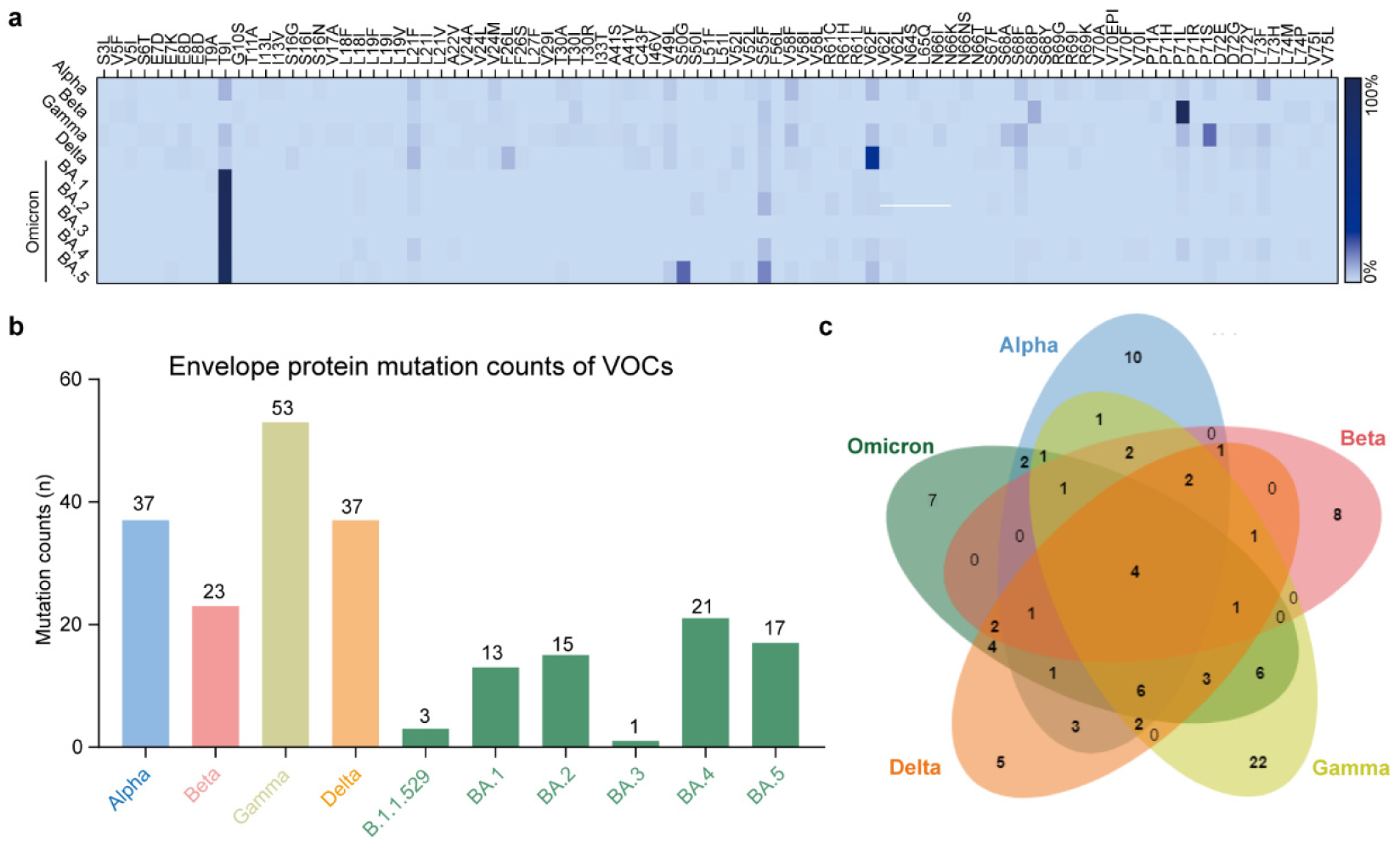
Statistics of 2-E mutations. **a**. Heat map of 2-E mutations frequency in five VOCs. **b**. The mutation counts of 2-E in each VOCs. **c**. Venn diagram of 2-E mutations in Omicron compared with other VOCs.

**Fig.S2.**
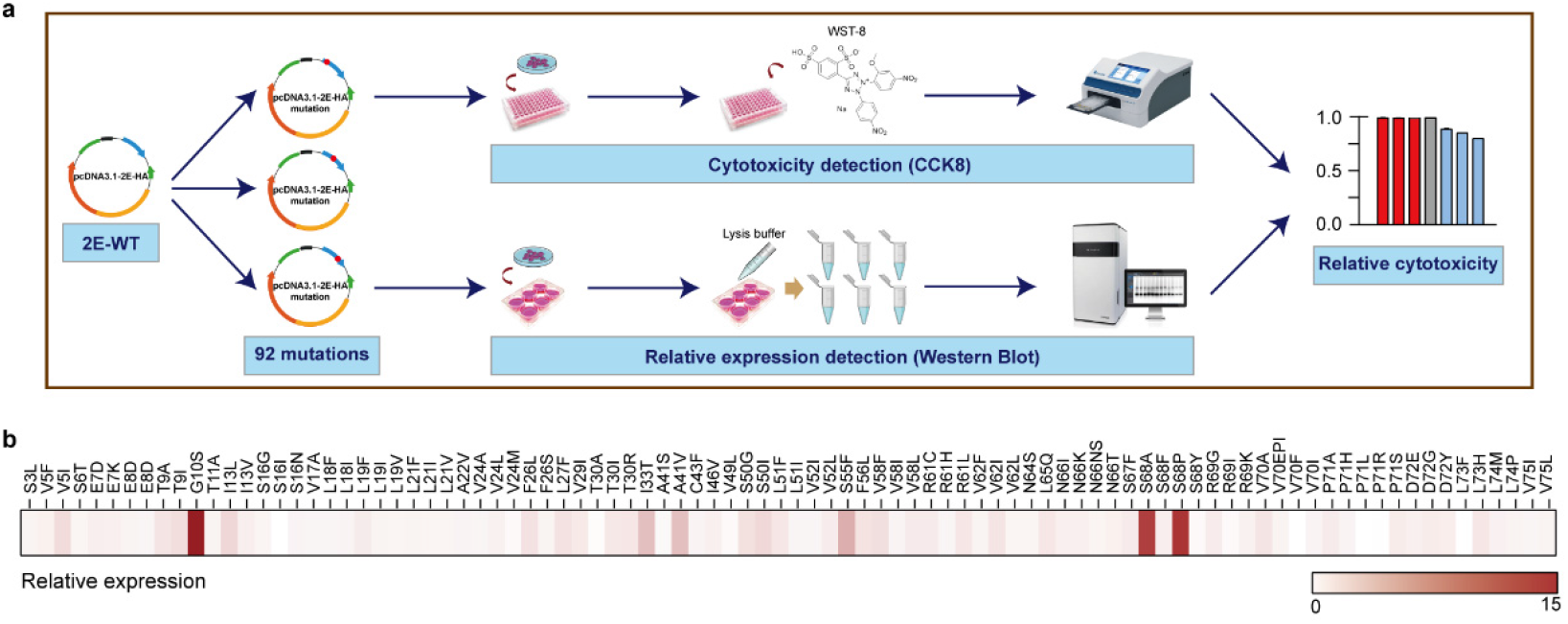
Cell lethality detection of 2-E mutations. **a**. Schematic of cell lethality detection. **b**. The protein expression level of 2-E mutations in Vero E6 cells after transfection.

